# Proteome-wide identification of metamorphic protein candidates using mass spectrometry

**DOI:** 10.1101/2025.11.12.688092

**Authors:** Baiyi Quan, Yanping Qiu, Kathleen Carillo, Andy LiWang, John Orban, Tsui-Fen Chou

## Abstract

How protein folds remodel at low temperature remains poorly mapped at the proteome scale. We combine brief, temperature-controlled proteolysis with LC–MS/MS to chart cold-dependent conformational changes in *Escherichia coli* lysate. Methodological validation with the metamorphic protein Sa1V90T—whose α/β-plait and 3α states interchange between 30 °C and 5 °C—showed that a high-protease, short-digestion regime detects temperature-dependent exposure at four peptides, whereas the temperature-insensitive variant Sa1V90TV52D is largely unchanged. Applied to lysate, the workflow identifies >750 proteolytic sites across >250 abundant proteins (>10% of the identified proteome) whose local susceptibility shifts as temperature decreases from 35 °C to 5 °C, including five Sa1V90T peptides recapitulating purified-protein behavior. A site-level scoring model yields 338 destabilized peptides (177 proteins) and 426 stabilized peptides (138 proteins). Cold stabilization prominently marks translation machinery (>15 ribosomal proteins), while cooling also exposes regions in chaperones and cell-division factors. Sequence analysis links temperature sensitivity to charge-enriched composition, consistent with thermodynamic expectations. These data provide a proteome-wide view of cold-sensitive folding, implicate translation and cytokinetic systems in low-temperature remodeling, and illuminate a new strategy for identifying metamorphic proteins on an unprecedented scale.

## Introduction

AI models like AlphaFold3^1^ and RoseTTAFold^2^ now predict static protein structures with accuracy, yet how folds remodel under different conditions remains poorly resolved. Temperature is a pervasive perturbation for protein folding. Unlike many ordered molecular systems, proteins often adopt maximal stability within a narrow thermal window and can denature at both high and low temperatures^3,4^. Compared with heat-induced unfolding, cold denaturation is less characterized because for most proteins the midpoint lies below the freezing point of aqueous buffers^3,5,6^. One driver of protein cold denaturation is the weakened hydrophobic effect at low temperature—a principal force stabilizing the folded state. Strategies such as *ad hoc* mutations^7–9^ or chemical denaturants^3,10^ can shift cold denaturation above 0 °C, and a few proteins (e.g., Yfh1, a yeast frataxin ortholog implicated in Friedreich’s ataxia^11,12^) have been shown to unfold near or above 0 °C. In addition, metamorphic proteins—polypeptides that switch between distinctly different native folds—have been hypothesized to bias toward alternative states at low temperature^13^. How these temperature sensitivities map onto cellular function remains largely unclear.

Unlike homeotherms, microbes cannot regulate body temperature and must accommodate ambient fluctuations. Understanding how bacteria tolerate or exploit cold has implications for food and drug preservation, environmental microbiology, and anti-bacterial strategies^14–16^. During *Escherichia coli* cold shock, cold-shock proteins (CSPs) are induced and hypothetically function as nucleic-acid chaperones, yet the upstream regulatory architecture is incompletely defined^15,17–19^. Over longer acclimation periods, bulk proteome composition can appear relatively stable across a limited temperature range^20,21^, even as growth rates follow Arrhenius-like scaling with absolute temperature—patterns often ascribed to enzyme-kinetic constraints^21–23^. Whether additional, biophysical mechanisms contribute—particularly below the Arrhenius regime—remains an open question.

Proteolysis-based readouts provide a sensitive, local measure of conformational exposure^24–26^. Coupled to LC–MS/MS, approaches such as Limited Proteolysis^27–30^ (LiP-MS) and the PEptide-centric Local Stability Assay^31^ (PELSA) enable proteome-scale detection of stability changes at peptide resolution. Here, we extend these methods to profile cold-dependent conformational remodeling in *E. coli* lysate. We map >750 proteolytic sites across >250 abundant in proteome proteins whose local susceptibility changes as temperature decreases—approximately an order of magnitude more protein-level observation than previously reported^17^. Many candidates intersect pathways implicated in cold adaptation or core bacterial stress response. These data provide a proteome-scale view of cold-sensitive folding states, infer novel mechanisms beyond those previously suggested, and inform the thermodynamics of protein stability in the cellular context. Importantly, they also report on potential metamorphic proteins, which have been challenging to identify both experimentally and computationally, and for which partial cold denaturation has been recently postulated to be a general feature^13^.

## Results

### In vitro validation of a proteolysis-based assay for temperature-sensitive folds

We developed a proteolysis-coupled proteomics workflow to detect temperature-dependent conformational changes (**Fig. 1A**). Proteins were equilibrated at defined temperatures and subjected to brief high-enzyme digestions (E:S 1:2–1:10). Prior work has shown that accelerated local cleavage can report on conformational exposure driven by interactions, ligand binding, or cellular state^27–32^. We reasoned that the same principle would capture temperature-induced changes, including cold sensitivity.

**Figure 1.**
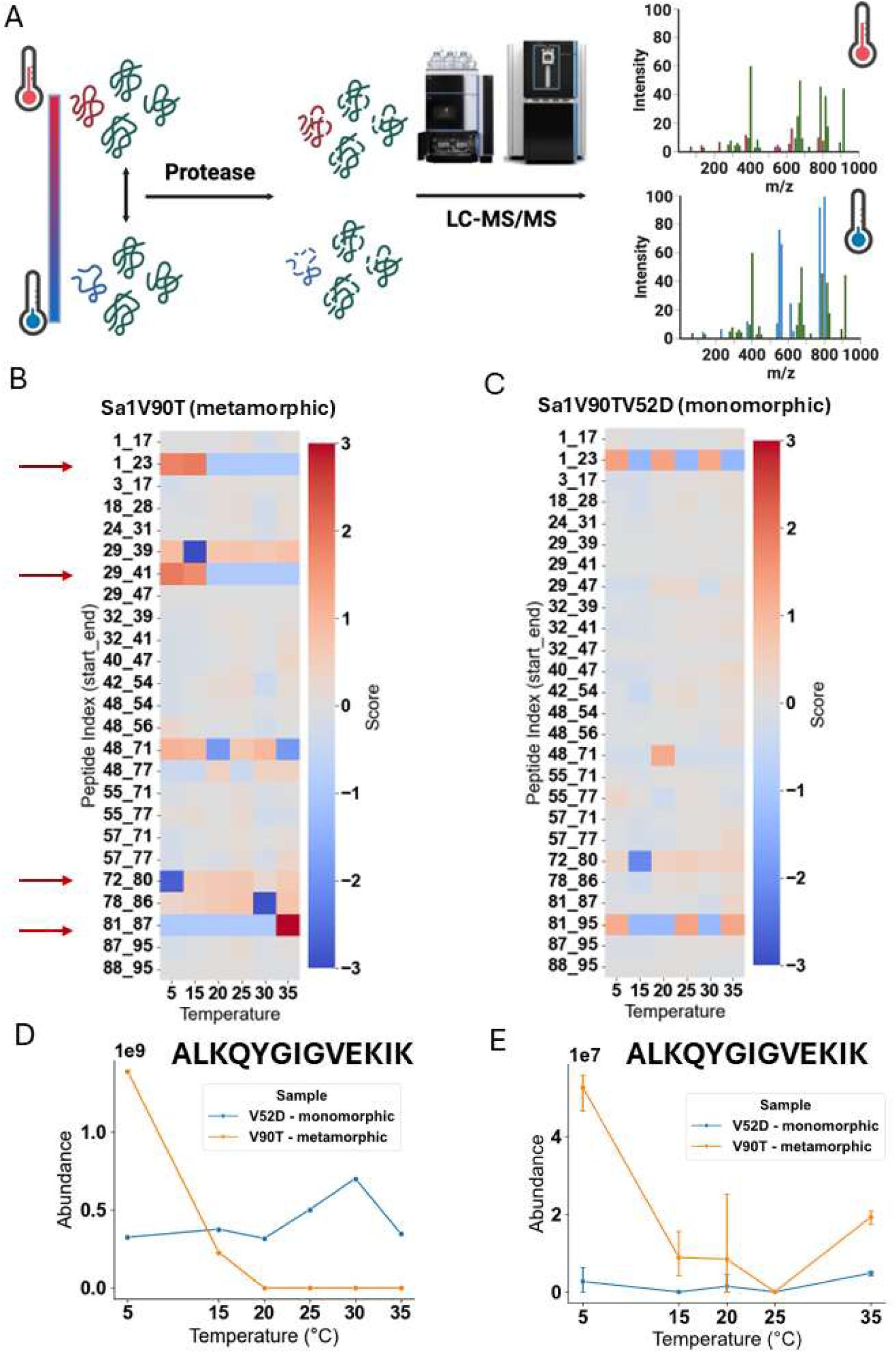
*In vitro* and *In vivo* validation of the method. **(A)** Proposed workflow of the study. Briefly, proteolysis happen under different temperatures. The proteins that change their folding states will have different exposed sites, hence different proteolysis patterns, which is detected by LC-MS/MS as a readout. **(B)** and **(C)** Heatmap showing the Sa1 peptides identified in the *in vitro* validation. The y-axis is the peptides, indexed by their starting and ending amino acid positions, and the x-axis is the temperature. The color stands for a score that is a weighted normalized fold change, which is given by score = I/I_avg x (I_maxP – I_min)/(I_max – I_min), where I is the intensity of the peptide under certain temperature. I_avg is the average intensity of the peptide across all temperatures, I_maxP is the maximum intensity of the peptide across all temperatures, and I_max and I_min is the maximum and minimum intensities of all the peptides identified for Sa1 across all temperatures, respectively. Peptides labeled with red arrows are the peptides whose intensities are temperature-dependent only in V90T (metamorphic) variant but not in V90TV52D (monomorphic). **(D)** and **(E)** Plots showing the abundance of one of the Sa1 temperature-dependent peptides, ALKQYGIGVEKIK (denoted as 29-41 in **(B)** and **(C)**), *in vitro* and *in vivo*, respectively. The y-axis is the abundance and x-axis is temperature. The color of the plot represents the variants.

As an in vitro test case, we examined the metamorphic protein Sa1V90T, which interconverts between two folds: an α/β-plait state predominates at 30 °C, a 3α state at 5 °C, and both are approximately equally populated near 17 °C^33^. A single-site variant, Sa1V90TV52D, resists this temperature switch and remains in the α/β-plait topology from 30 °C to 5 °C. To minimize temperature-dependent changes in protease activity and reduce oligomerization at the working concentrations, purified Sa1V90T and Sa1V90TV52D were spiked into bovine serum albumin (BSA). BSA peptide intensities were used for activity normalization.

We compared two digestion regimes: a shorter, high-trypsin protocol^31^ (**Fig. 1B**-**C**, **Tables S3-4**, **S9**-**10**) and a longer, lower-trypsin protocol (**Supple. Fig. 2**, **Tables S5-6**, **S11**-**12**), as well as limited proteolysis protocol using proteinase K^28^ (**Supple. Fig. 1**, **Tables S1-2**, **S7**-**8**). The high-trypsin regime—describe as PELSA (PEptide-centric Local Stability Assay) by Li et. al.^31^—more sensitively captured temperature-dependent behavior in Sa1V90T while remaining largely unchanged for Sa1V90TV52D (**Figs. 1B-D**). Four Sa1V90T peptides exhibited temperature-dependent proteolysis (**Supple. Fig. 3**). Three reported cold-induced destabilization (partial cold denaturation), consistent with prior NMR observations. These data demonstrate that the workflow resolves fold-state changes driven by temperature.

**Figure 2.**
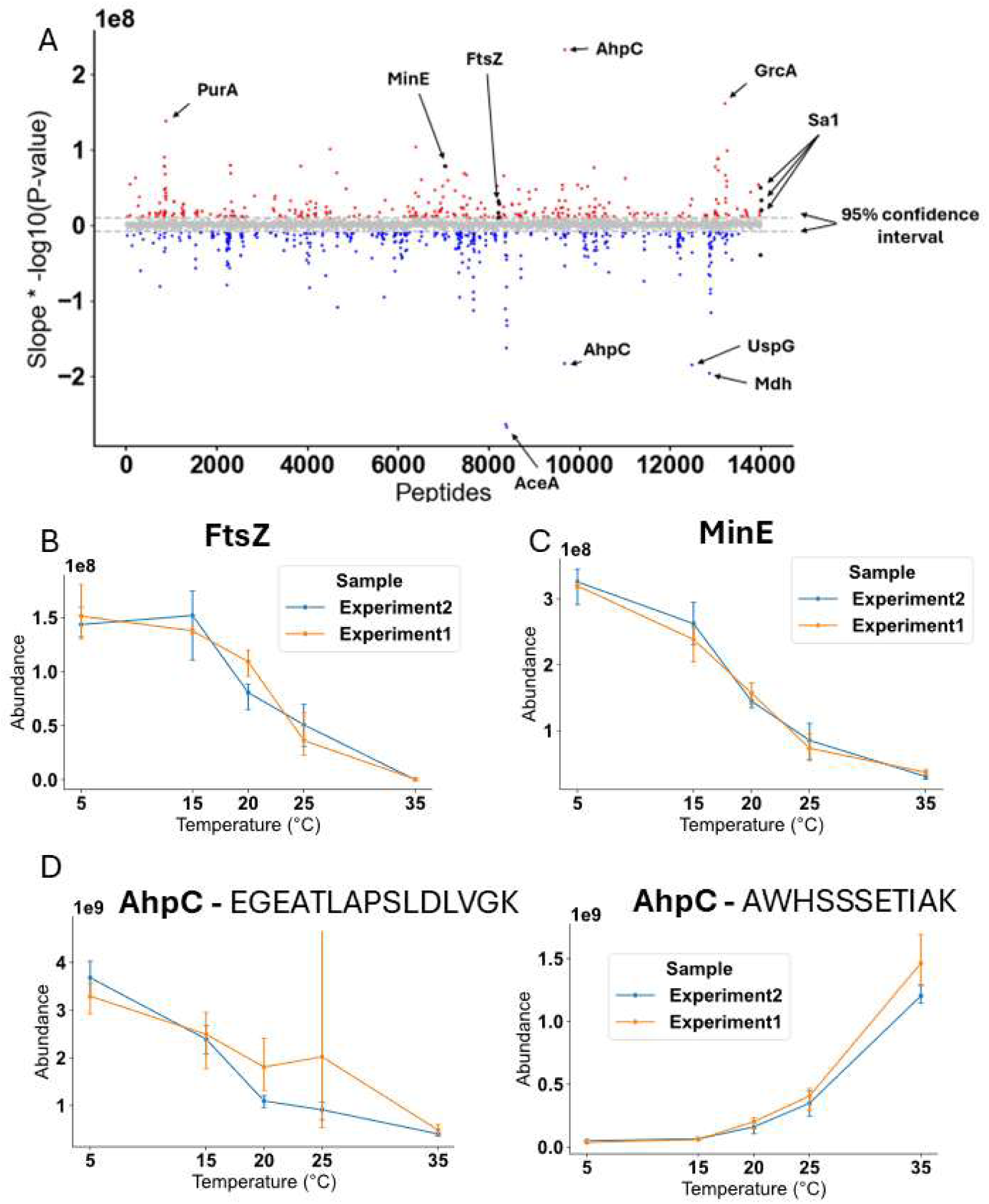
Global Profiling of Cold-stabilized and Cold-destabilized Proteins. **(A)** Manhattan plot of the analysis. Each dots represents one peptide. The x-axis is the peptides (grouped by proteins), and the y-axis is the slope x -log10(P-value) where both slope and p-value are calculated by fitting the abundances and the temperature into a linear model. Red dots represents destabilized peptides, and blue dots represents stabilized peptides. Gray dots represents the peptides with no significant temperature dependency, and the gray horizontal lines stands for 95% confidence interval from fitting the distribution into a Student’s t-distribution. Peptides of known metamorphic proteins (Sa1, FtsZ, and MinE) are highlighted with black color. **(B), (C), (D),** and **(E)** Selected examples of peptides from two metamorphic proteins (FtsZ and MinE), and two peptides from AhpC, which shows different temperature dependencies, respectively. The y-axis is the abundance and x-axis is temperature. The color of the plot represents two experiments, each with three replicates.

**Figure 3.**
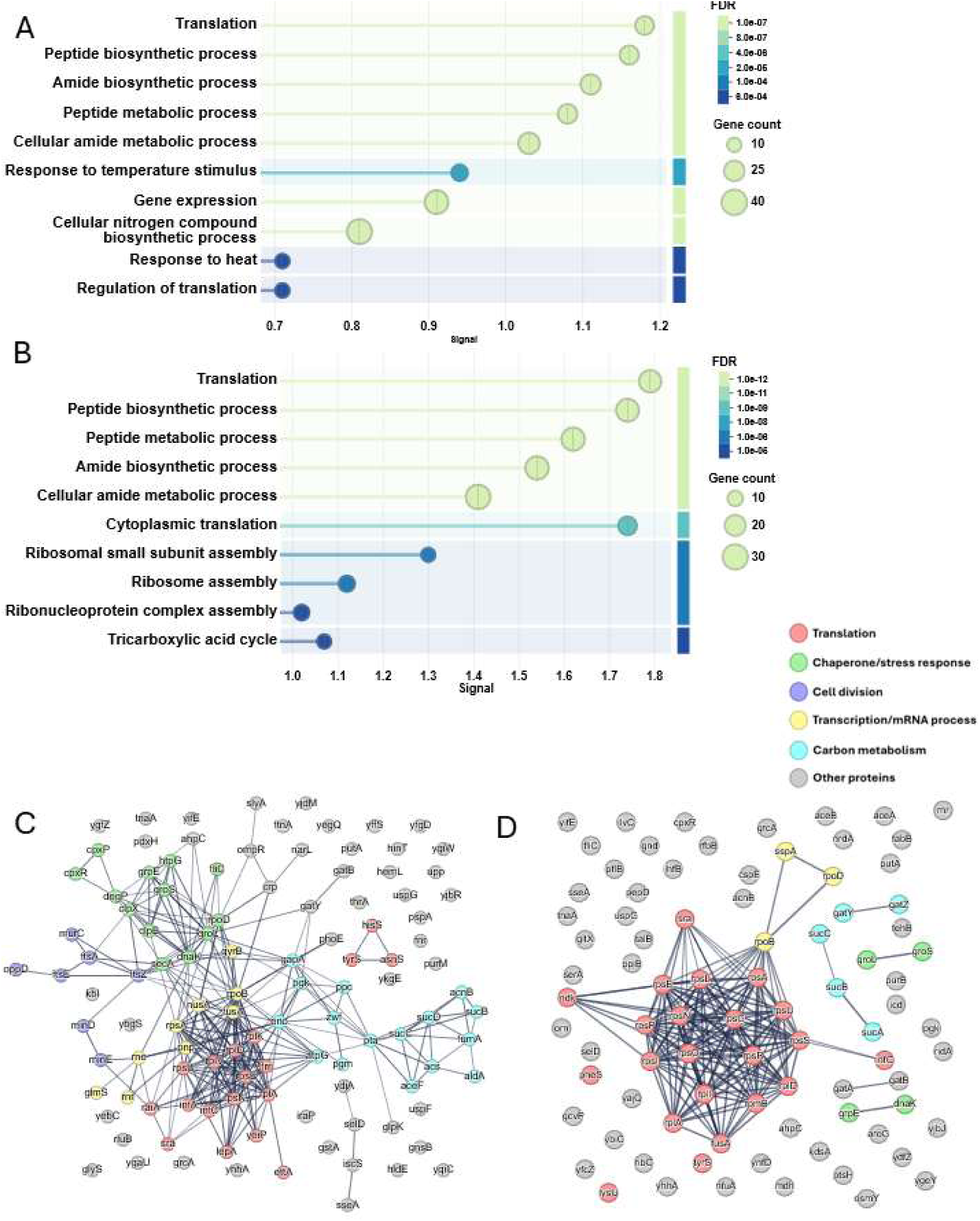
Functional and Interactome Analysis of the Cold-stabilized and Cold-destabilized Proteins. **(A)** and **(B)** Gene ontology (GO) over-representation analysis (biological process) of the cold-stabilized and cold-destabilized proteins, respectively. The y-axis is the biological process terms from Gene Ontology database, and the x-axis is the signal, which is defined by a weighted harmonic mean between the observed/expected ratio and -log(FDR). The color represents FDR and the size of the circles represents gene counts of the term. **(C)** and **(D)** Interactome of the cold-stabilized and cold-destabilized proteins, respectively. Only physical interaction is shown in the plots. The weight of the lines between dots represents how strong the evidence of the interaction is. The proteins are colored by their biological function.

### Proteome-wide profiling in lysate

We next asked whether the behavior persists in complex mixtures. Sa1V90T and Sa1V90TV52D were spiked into *E. coli* lysate and analyzed by LC–MS/MS (**Tables S13**-**14**). Five Sa1V90T peptides showed temperature-dependent proteolysis in lysate, including the four observed with purified protein. As temperature decreased from 35 °C to 5 °C, four peptides were destabilized (increased susceptibility) and one was stabilized, mirroring the purified-protein analysis (**Fig. 1E**).

We then assessed endogenous proteins detected in the lysate. For each peptide, intensity was regressed against temperature in a simple linear model. We scored peptides as score = slope × log10(p), where p is a p-value calculated from a two-sided t-test for the null hypothesis that the true slope is 0. Peptides with increasing intensity at lower temperature (destabilized/denatured upon cooling) yield positive scores, whereas those with decreasing intensity (stabilized/protected upon cooling) yield negative scores. The score distribution was modeled with a Student’s t distribution (**Supple. Fig. S5**) to derive 95% confidence interval (CI). Peptides outside the CI were called “hits”. This yielded 338 destabilized peptides (177 proteins) and 426 stabilized peptides (138 proteins) (**Figs. 2A**, **Table S15**). The overall peptide-level hit rate was ∼5.5% (764/13994), close to the nominal two-tailed 5% threshold, indicating that orthogonal validation will be important for refining specificity. Notably, 51 proteins harbored sites that were both cold-stabilized and cold-destabilized, including AhpC (**Figs. 2D-E**), indicating that cold responses are region-specific rather than uniformly protective or destabilizing.

### Biological processes associated with cold-sensitive folding

Gene-set inspection highlighted distinct trends. Among cold-stabilized proteins, translation components were prominent, including >15 ribosomal subunits, multiple aminoacyl-tRNA synthetases, and other ribosome-associated factors (**Figs. 3A**, **C**). While proteomics is biased toward abundant proteins, these data suggest that *E. coli* ribosome structure/assembly is reconfigured at low temperature. Among cold-destabilized proteins, we observed (albeit more modestly) additional translation factors (7 ribosomal subunits), heat-response proteins (e.g., chaperones), transcriptional machinery, and cell-division components (**Figs. 3B, D**). Notably, several chaperones and co-chaperones were destabilized on cooling, hinting that *E. coli* may mitigate cold stress via mechanisms distinct from classical heat-shock responses.

Cell-division proteins were enriched among hits, including multiple filamenting temperature-sensitive (Fts) factors. Historically, fts mutants divide at 30 °C but fail at 42 °C, implying intrinsic temperature sensitivity^34^. FtsA^35^ (actin-like) and FtsZ^35,36^ (tubulin-like), together with MinE of the MinCDE positioning system^37–39^ that regulates the FtsZ ring, featured among candidates (**Figs. 2B-C**). Both FtsZ and MinE have been proposed as metamorphic proteins, analogous to Sa1^40^.

### Sequence determinants of temperature sensitivity

To probe sequence-encoded features, we computed 42 physicochemical descriptors for the *E. coli* proteome and compared cold-destabilized and cold-stabilized proteins to the identified background set using existing tools^41^ (**Figs. 4A-B**). Temperature-sensitive sequences were enriched for charge-related properties (e.g., lysine and glutamate content, fraction of charged residues, mean net charge). Cold-destabilized proteins tend to be enriched in positively charged residues. Cold-stabilized proteins showed increased positive and negative charges, yielding near-neutral mean net charge relative to background (**Figs. 4C-E**). These observations align with expectations from folding thermodynamics: the temperature dependence of folding free energy is shaped by heat capacity changes, which correlate with accessible surface area and, by extension, with polar/charged exposure^3,5,6,42^.

**Figure 4.**
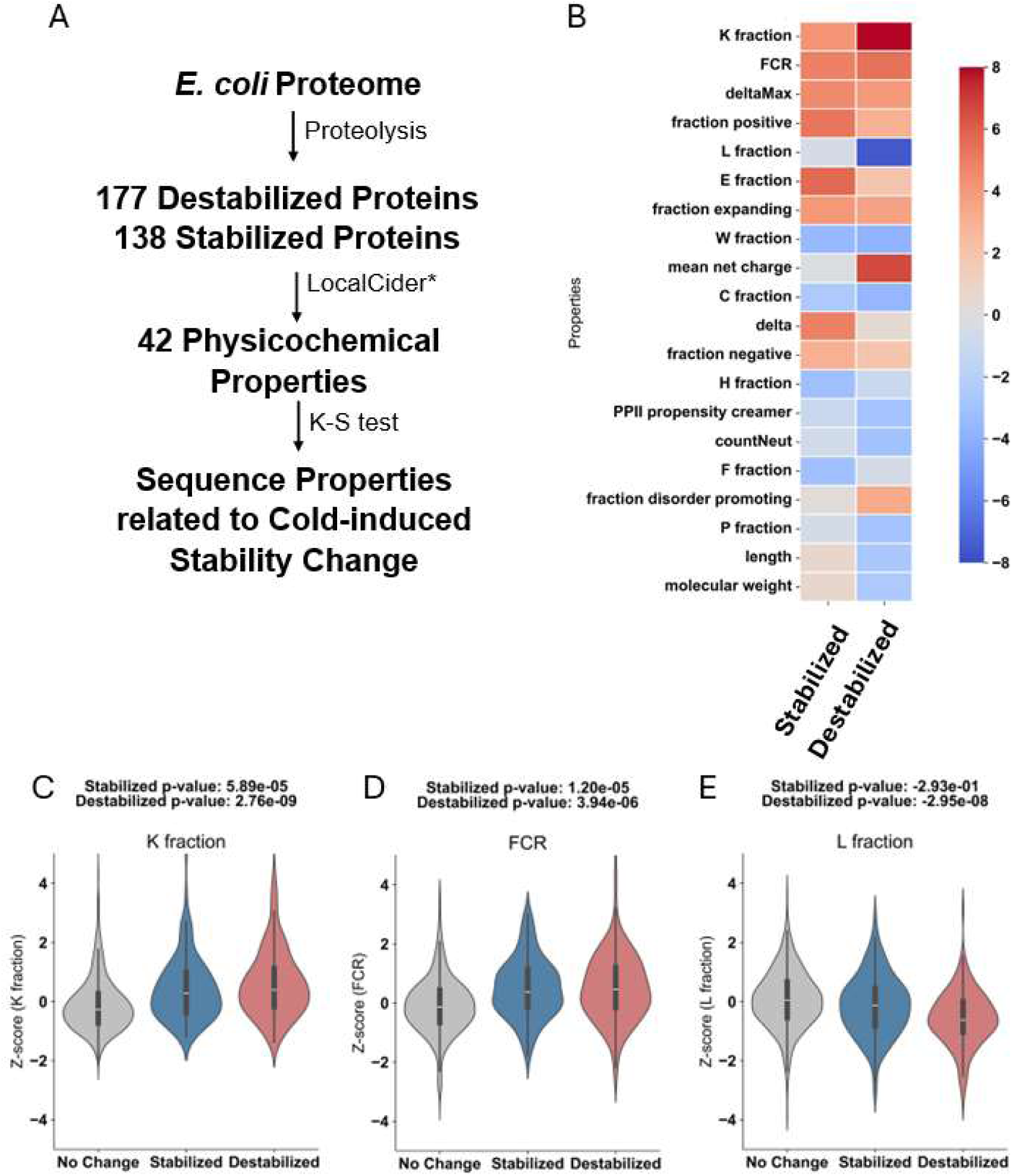
Sequence Properties of the Cold-stabilized and Cold-destabilized Proteins. **(A)** Workflow of the analysis. 42 physicochemical properties for the E. coli proteome are calculated by LocalCider, and Kolmogorov-Smirnov test is used to calculate the p values to find out the sequence properties for related to cold-induced stability change. **(B)** Heatmap showing properties related to cold-stabilized and cold-destabilized proteins. The x-axis is the cold-induced behavior, and the y-axis is the properties. The color represents the signed p-values calculated by Kolmogorov-Smirnov test. **(C), (D),** and **(E)** Violin plots for examples of properties. The x-axis (and the color) is the cold-induced behavior, and the y-axis is the Z-score of the property.

## Discussion

Cold denaturation is well documented thermodynamically, yet systematic identification of proteins that partially unfold above 0 °C has been limited. Traditional approaches (e.g., Differential Scanning Calorimetry) often require additional perturbations (pH/denaturant) to shift midpoints above the freezing point, complicating proteome-scale surveys^3,10^. Here, a brief proteolysis readout coupled to LC–MS/MS detects both cold-induced exposure and protection at individual proteolytic sites, enabling candidates that denature—or become conformationally protected—upon cooling in aqueous buffer.

The “cold stabilization” phenomenon observed at specific sites likely reflects allosteric or quaternary-structure changes, consistent with proteins that display both protected and exposed regions in the same polypeptide. Non-physiological oligomerization in lysate could contribute in some cases. A subset of hits overlaps with proteins predicted by IUPred3^43^ to contain >30-residue intrinsically disordered regions (IDRs). Although the overall IDR proportion is similar to the *E. coli* proteome background, prior reports suggest certain IDRs adopt structure at low temperature^44,45^. Targeted validation will be needed to resolve these possibilities and to probe their roles in cold adaptation.

The dataset also reveals putative metamorphic proteins (e.g., FtsZ, MinE). Discovery of metamorphic proteins has been largely serendipitous and relies on time- and material-consuming approaches^13,40^ (NMR, SEC, etc.). A proteolysis-based, site-resolved proteomic screen offers a scalable and more high-throughput complementary approach. The principal limitation is specificity: with thresholds set by distributional modeling, expected false positives remain. However, integration with emerging AI models for fold heterogeneity (e.g., CF-Random^46^, AF-Cluster^47^) and iterative validation should improve prioritization while providing much-needed training data for these predictors.

A principal limitation of our approach is reduced proteome coverage arising from the non-canonical, high-enzyme/short-duration digestions. These conditions were chosen to enhance sensitivity and site-level specificity for detecting cold-stabilized or cold-destabilized regions, but they concomitantly curtail sequence sampling and hinder identification of low-abundance proteins. Additional optimization is needed to balance specificity with coverage—for example, by tuning enzyme dose and digestion time, incorporating orthogonal proteases, or pairing with deeper LC–MS/MS acquisition. Consistent with this trade-of, FtsH and IscA—whose orthologs have been reported as metamorphic in other organisms^40^—were detected with partial cold-destabilizing behavior but not called as hits (**Supple. Fig. 4**), likely due to low abundance. Other known metamorphic proteins (e.g., RfaH and cytotoxin^40^) were not detected, presumably reflecting limited coverage under the current conditions.

## Conclusions

Across the *E. coli* lysate, we map >750 proteolytic sites on >250 proteins whose local proteolytic susceptibility increases as temperature is lowered from near-physiological to 5 °C—constituting, to our knowledge, the largest compendium of cold-sensitive proteolytic readouts to date. The results (i) reinforce thermodynamic expectations at the proteome level, (ii) implicate translation and cell-division systems in low-temperature remodeling and cold adaptation, and (iii) highlight candidates for metamorphic behavior. More broadly, our approach provides a practical entry point for charting temperature-sensitive conformational landscapes and for screening for proteins whose folds—and functions—are rewired by cold.

## Acknowledgement

The Proteome Exploration Laboratory is partially supported by Caltech Beckman Institute Endowment Funds. We also acknowledge support from US NIH grants R35GM144110 (to A.L.) and R01GM062154 (to J.O.), US Army grant W911NF-23-1-0248 (to A.L.), and NSF-CREST: Center for Cellular and Biomolecular Machines at the University of California, Merced (NSF-HRD-1547848).

## Author contributions

Conceptualization: B.Q., A.L., J.O., and T.F.C. Funding acquisition: T.F.C. Experiment: B.Q., Y.Q., and K.C. Data analysis: B.Q. Supervision: A.L., J.O., and T.F.C. Writing—original draft: B.Q. Writing—review and editing: A.L., J.O., and T.F.C.

---

**Supplementary Figure S1.**
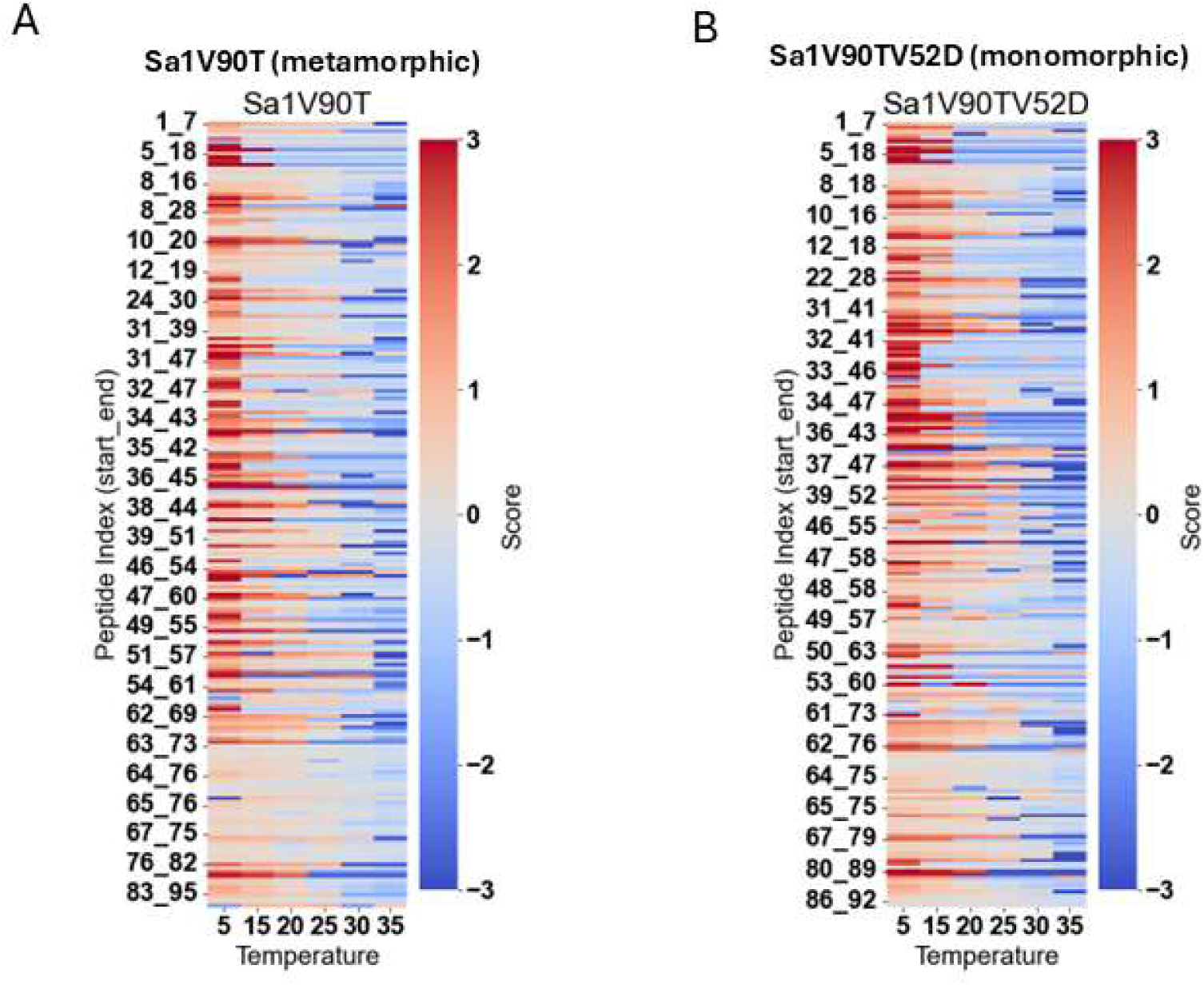
Limited Proteolysis under Different Temperature *in vitro*. **(A)** and **(B)** Heatmap showing the Sa1 peptides identified in the *in vitro* limited proteolysis experiment. The y-axis is the peptides, indexed by their starting and ending amino acid positions, and the x-axis is the temperature. The color stands for a score that is a weighted normalized fold change, which is given by score = I/I_avg x (I_maxP – I_min)/(I_max – I_min), where I is the intensity of the peptide under certain temperature. I_avg is the average intensity of the peptide across all temperatures, I_maxP is the maximum intensity of the peptide across all temperatures, and I_max and I_min is the maximum and minimum intensities of all the peptides identified for Sa1 across all temperatures, respectively. The abundances are normalized using the BSA peptide abundances. No significant changes are observed between metamorphic and monomorphic variants.

**Supplementary Figure S2.**
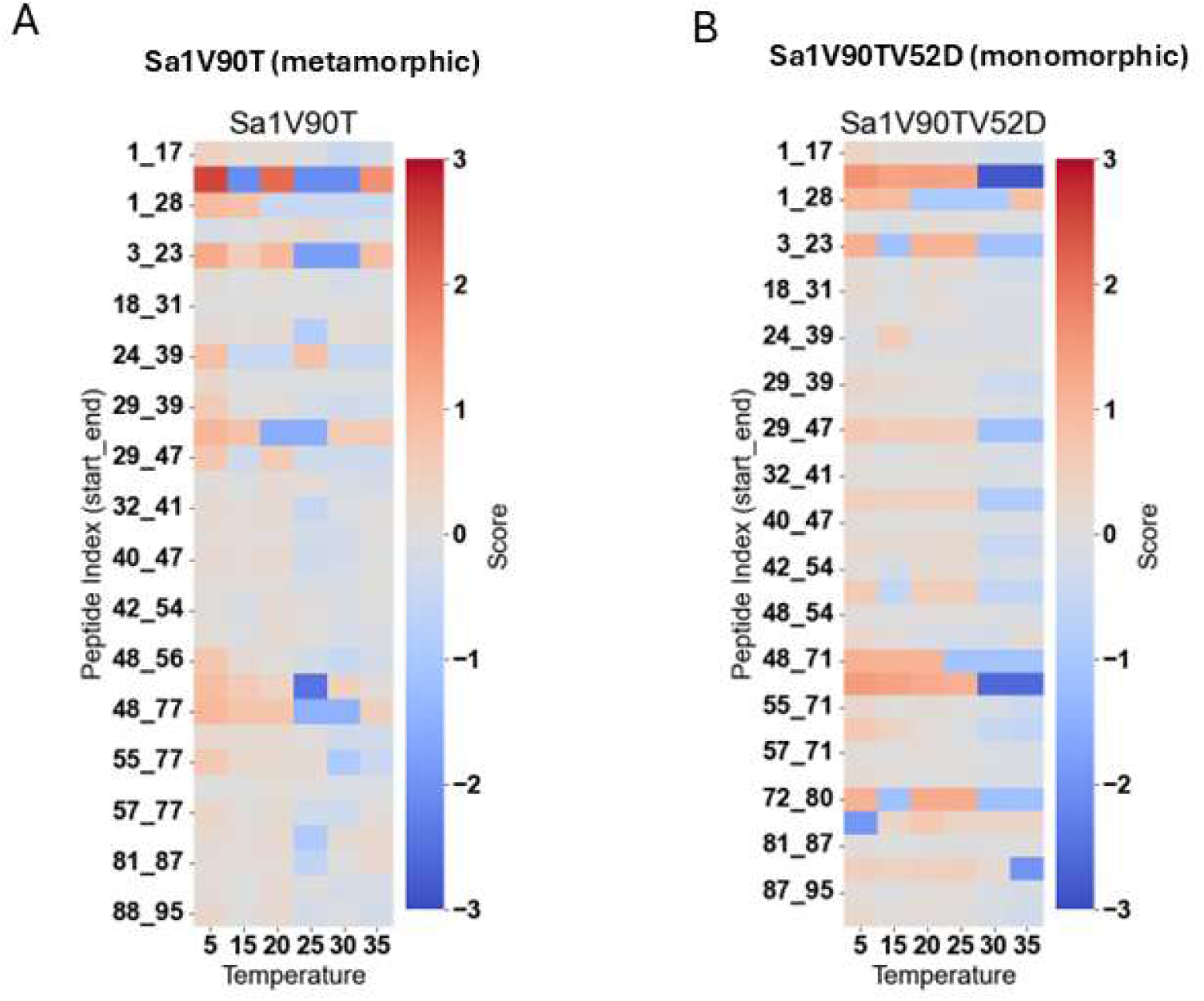
Low-Trypsin Proteolysis under Different Temperature *in vitro*. **(A)** and **(B)** Heatmap showing the Sa1 peptides identified in the *in vitro* low-trypsin regime experiment. The y-axis is the peptides, indexed by their starting and ending amino acid positions, and the x-axis is the temperature. The color stands for a score that is a weighted normalized fold change, which is given by score = I/I_avg x (I_maxP – I_min)/(I_max – I_min), where I is the intensity of the peptide under certain temperature. I_avg is the average intensity of the peptide across all temperatures, I_maxP is the maximum intensity of the peptide across all temperatures, and I_max and I_min is the maximum and minimum intensities of all the peptides identified for Sa1 across all temperatures, respectively. The abundances are normalized using the BSA peptide abundances. No significant changes are observed between metamorphic and monomorphic variants.

**Supplementary Figure S3.**
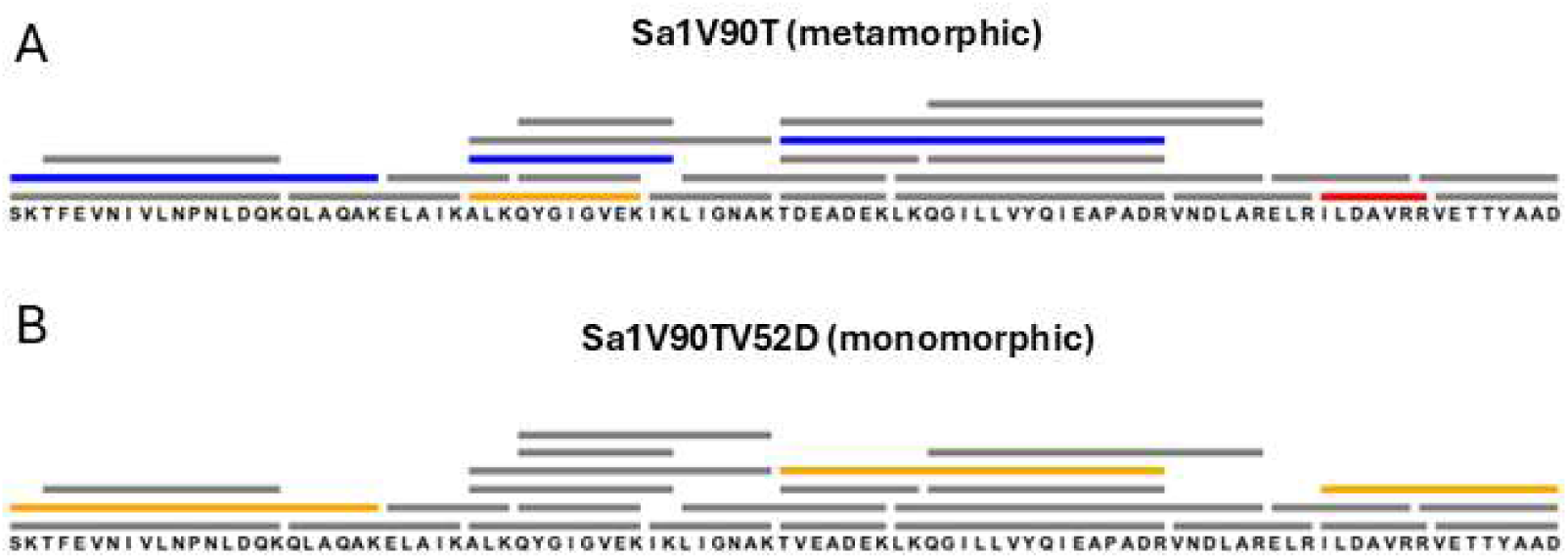
Low-Trypsin Proteolysis under Different Temperature *in vitro*. **(A)** and **(B)** Sequence map showing the localization of Sa1 peptides identified in the *in vitro* high-trypsin regime (PELSA) experiment for V90T and V90TV52D variant, respectively. The blue bars indicate the peptides that are destabilized as temperature decreases, red bars indicate the peptides that are stabilized as temperature decreases, and the gray bars indicate the peptides that are not significantly changed as temperature changes. The yellow bars indicate the peptides whose abundances varies drastically as temperature changes but no significant trends can be found.

**Supplementary Figure S4.**
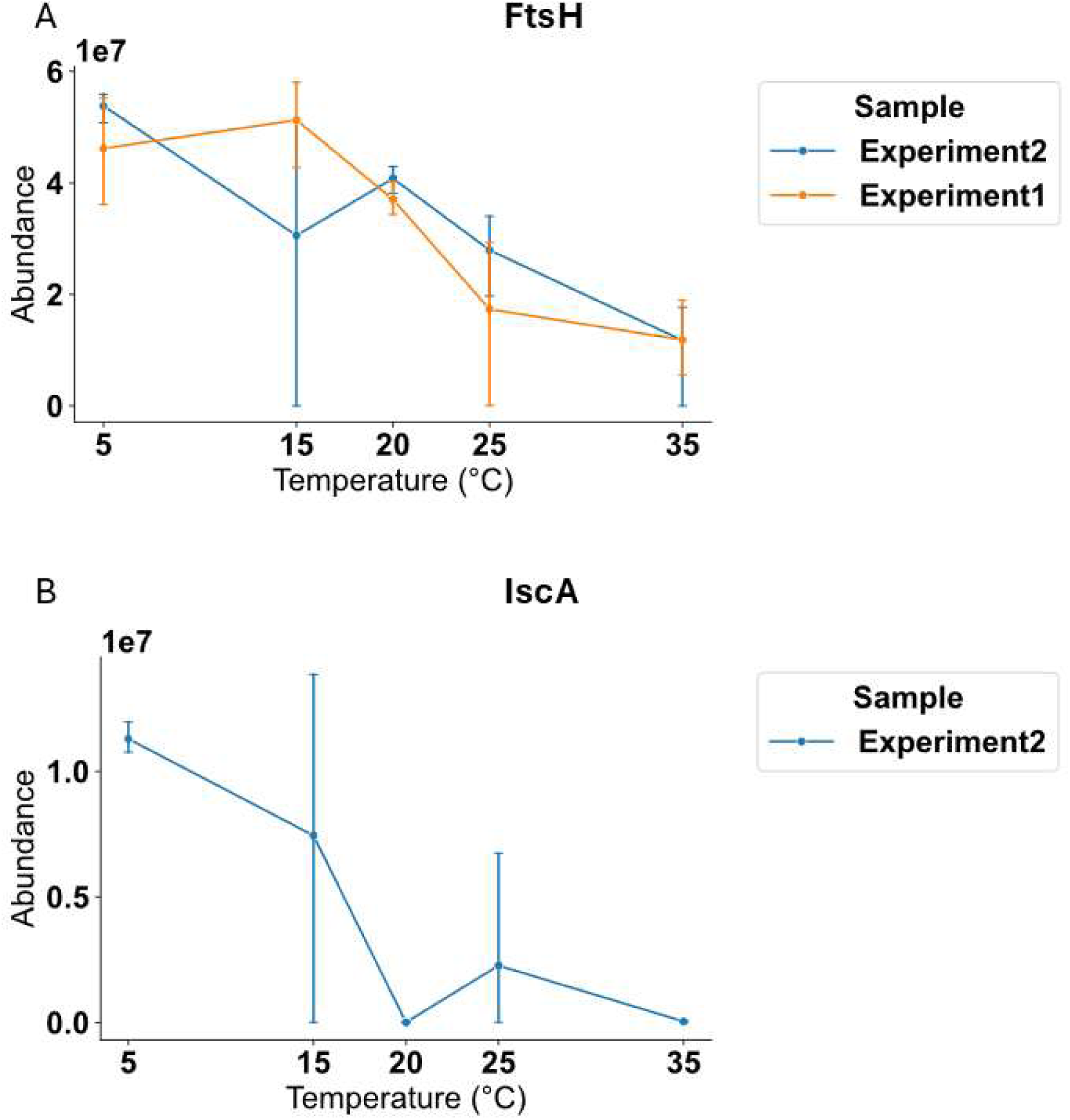
More Representative Peptides from Potential Metamorphic Proteins. **(A)** and **(B)** Selected examples of peptides from two potential metamorphic proteins that was not a hit in our analysis (FtsH and IscA). The y-axis is the abundance and x-axis is temperature. The color of the plot represents two experiments, each with three replicates.

**Supplementary Figure S5.**
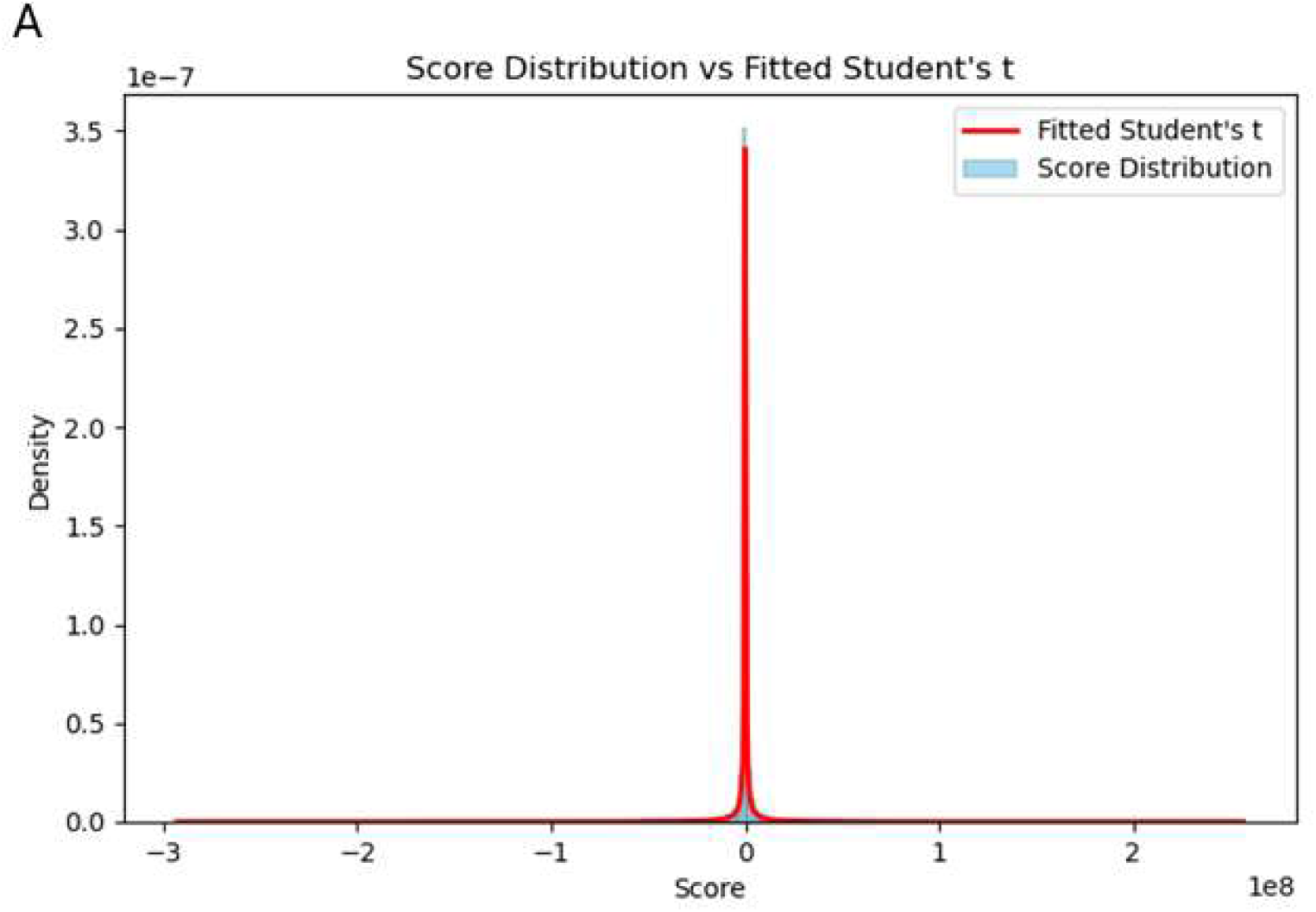
T-distribution Fitting for the Slope x –log10(p value). **(A)** Fitting the score = Slope x –log10(p value) into a Student’s t-distribution shows good fitting.

