## Supplementary Methods for "Proteome-wide identification of metamorphic protein candidates using mass spectrometry"

**Expression and Purification of Sa1 Variants**

Site-directed mutants were prepared using mutagenic primers and a Q5 site-directed mutagenesis kit (BioLabs). The Sa1 and its variants were then cloned into the eXact tag pH0720 vector, which contains an engineered subtilisin prodomain and an HSA-binding GA sequence as N-terminal tags. These tags were removed during purification. BL21(DE3) strain E. coli cells were transformed with the vector and cultured in LB media. The cells were incubated at 37 °C until they reached an OD600 of 0.6 to 0.8. Protein expression was induced by adding 1 mM IPTG and incubating the cells at 25 °C for 18 hours. After incubation, the cells were harvested by centrifugation and resuspended in a buffer containing 100 mM potassium phosphate (KPi) at pH 7.0, with 150 mM NaCl and a protease inhibitor cocktail tablet (Pierce). The cells were then lysed using sonication. The soluble cell extract containing the prodomain-GA fusion protein was loaded onto a 5 mL subtilisin column. The column was washed with five column volumes of 100 mM KPi (pH 7.0), followed by 20 column volumes of 500 mM NaCl in 100 mM KPi (pH 7.0) to remove impurities. After washing with a high-salt-concentration buffer, an additional wash with 5 column volumes of 100 mM KPi was performed. To cleave and elute the purified protein, 6 mL of a 3.5 mM imidazole solution in 100 mM KPi (pH 7.0) was injected at a flow rate of 1 mL/min. The eluent was then passed through an HSA column to eliminate any uncleaved fusion protein, and the cleaved target protein was collected in the flow-through. Fractions containing high-purity protein, as determined by SDS-polyacrylamide gel electrophoresis, were pooled and concentrated for further analysis.

**Proteolysis Experiment to Measure Protein Cold Stability**

Purified Sa1 variants were spike-in to BSA solution or *E. coli* lysates with a spike-in concentration of 0.05 – 0.3 mg/ml, and total protein concentration after spike-in of around 1 mg/ml. The samples aliquoted into 20 µl aliquots for proteolysis reaction, and were equilibrated at corresponding temperatures for more than 10 min before proteolysis. For high-trypsin proteolysis (PELSA method^31^), trypsin-TPCK (Thermo) was added into the sample with an enzyme:substrate ratio of 1:2, and the proteolysis was allowed for 1 min at certain temperature before quenched with final concentration of 50 mM PMSF. For low-trypsin proteolysis, trypsin-TPCK (Thermo) was added into the sample with an enzyme:substrate ratio of 1:10, and the proteolysis was allowed for 10 min at certain temperature before quenched with final concentration of 50 mM PMSF. For proteinase K proteolysis (LiP method^28^), proteinase K from *Tritirachium album* (Sigma-aldrich) was added into the sample with an enzyme:substrate ratio of 1:100, and the proteolysis was allowed for 5 min at certain temperature before quenched with final concentration of 50 mM PMSF. The samples were later acidified using 200 µl of 1% TFA and desalted using Peptide Clean-up Plate (Thermo) according to manufacturer’s protocol. The desalted samples were dried using Refrigerated CentriVap Concentrator (LabConco) and stored at -80 °C before LC-MS/MS analysis.

**LC-MS/MS analysis**

The sample in the autosampler vial was dissolved in 20 µl of buffer A (2% acetonitrile, and 0.2% formic acid in water). The vial was briefly centrifuged and transferred to the autosampler. The LC-MS/MS analysis was performed using a Thermo Astral Mass Spectrometer coupled with a Thermo Vanquish Neo UHPLC system. The samples were analyzed on an Aurora Ultimate TS25 C18 column (25 cm x 75 µm, Ionopticks) coupled with a PepMap™ Neo Trap Cartridge (300 µm x 5 mm, Thermo). Buffer A was 2% acetonitrile, and 0.2% formic acid in water, and Buffer B was 80% acetonitrile, and 0.2% formic acid in water. For BSA spike-in runs, the gradient was 2-6% B for 2 min, 6-25% B for 10 min, 25-40% B for 3 min, 40-98% B for 1 min, and 98% B for 4 min. For *E. coli* lysate spike-in runs, the gradient was 2-5% B for 1 min, 5% B for 0.1 min, 5-35% B for 60 min, 35-70% B for 2 min, 70-99% B for 4 min, and 99% B for 4.9 min. The flow rate was 350 nl/min for both gradients. The spray voltage was 1900V in positive mode, and the ion transfer tube temperature was kept at 300°C. The full scan was set at 120000 resolution with a scan range of 375-1500 Th. The RF lens was set at 45%. The AGC target was set at 3.0 E6 at 300%, and the max ion injection time is 100 ms. Only peptides with charge state 2-5 was selected for MS2 acquisition, and the dynamic exclusion was set at 15 s with 5 ppm mass tolerance on both sides. The HCD collision energy was set to 29 NCE. The MS2 scan range is 110-1500 Th. The MS2 AGC target was set at 3.0 E3 at 30%, and the MS2 max ion injection time was 100 ms. The DDA cycle time was set at 0.6 seconds.

**Proteomics Data Analysis**

The raw data was searched using Proteome Discoverer 3.1 (Thermo) using SEQUEST search engine. Since the *E. coli* proteome retrieved from UniProtKB did not contain some of the reported metamorphic protein sequences, metamorphic protein sequences retrieved from Lauren et. al.^40^ was added to the database as well as Sa1V90T or Sa1V90TV52D sequences for corresponding spike-in samples. For trypsin samples, the enzyme was set as trypsin(full), and 4 missed cleavages were allowed for the search. For proteinase K samples, the enzyme was set as unspecific, and 3 missed cleavages were allowed for the search. For both enzymes, oxidation (+15.995 Da) on methionine was included as dynamic modifications. The search result was exported from Proteome Discoverer as .xlsx files and was processed using Python 3.12.7. The BSA peptides/total peptides abundances were used to normalize the enzyme efficiency at different temperatures. After normalization, the peptide abundances were later fit to a linear model with regards to the temperature using Scipy 1.13.1. The plots were generated using python packages including Matplotlib 3.9.2, Seaborn 0.13.2, and Pandas 2.2.2.

**IDR Sequence Prediction**

IDR sequences were predicted using IUpred3^43^ on *E. coli* proteome (acquired from UniProtKB). The cutoff of the iupred3 score was 0.5, and all regions that have more than 3 amino acids that have higher score than 0.5 were listed. The nearby regions were merged if they are apart from each other by less than 10 amino acids. Any regions shorter than 30 amino acids were removed, and only IDR sequences that are longer than or equal to 30 amino acids were subjected to further analyses.

**Extraction of the Features of the *E. coli* Proteins**

42 Physicochemical properties of *E. coli* proteins were calculated using localCIDER^41^ v0.1.20. The statistical analysis was performed using python 3.12.7 with packages including scikit-learn 1.5.1 and scipy 1.13.1. Briefly, all parameters calculated were Z-score normalized according the following equation:

$$Z= \frac{para-mean(para)}{std(para)}$$

where para stands for the parameter. Proteins that were cold-stabilized or cold-destabilized were extracted, and the selected proteins were compared to all the identified proteins in the proteomic dataset as a background. The comparison was performed per parameter and p value was reported using a Kolmogorov-Smirnov test to extract the features of interest. The reported signed p-value were calculated by following equation:

$$signed p=Sgn\left( {mean\left( para \right)}_{poi}-{mean\left( para \right)}_{background} \right)*(-{log}_{10}p)$$

where para stands for parameter.
